# Metabolic Stress Accelerates Dysregulated Synovial Macrophage-Fibroblast Communication and Htra1 Overproduction in Osteoarthritis

**DOI:** 10.1101/2024.09.26.615231

**Authors:** Garth Blackler, Joseph Klapak, Qinli Guo, Holly T. Philpott, HanYu Jiang, Dariana Ocica, Luigi Del Sordo, Benoit Fiset, Logan A. Walsh, C. Thomas Appleton WOREO Knee Study Group

## Abstract

Biomechanical and metabolic factors increase the risk for osteoarthritis (OA) by causing supraphysiological stresses on joint tissues. Chronic exposure to these stresses contributes to failure of the joint organ system, resulting in pain and loss of function for patients with OA. The synovium is vital for joint organ health but during OA, synovial inflammation and damage are associated with worse outcomes including pain. Unfortunately, the separate and combined effects of metabolic and biomechanical stresses on synovial tissues are not well understood. In this study, metabolic syndrome (MetS) was associated with worse knee pain in patients with early-stage knee OA, suggesting that metabolic stress may act on synovial tissues during early-stage OA, exacerbating outcomes. In a rat model of experimental knee OA, the combined effects of biomechanical and metabolic stresses induced worse knee pain, cartilage damage, and synovial inflammation than biomechanical stress alone. Further, single-cell RNA sequencing of synovial macrophages and fibroblasts identified earlier metabolic (glycolytic and respiratory) shifts, neurogenesis, dysregulated communication, and cell activation when metabolic and biomechanical stresses were combined. Lastly, using a direct contact co-culture system, we showed that metabolic stress alters macrophage-fibroblast communication leading to increased expression of Htra1, a pathogenic protease in OA. This study identifies novel mechanisms that may represent amenable therapeutic targets for patients experiencing MetS and OA.

**One-sentence summary:** Metabolic stress may cause worse outcomes in OA through dysregulated synovial cell communication that activates synovial fibroblasts and increases Htra1 production.

## Introduction

Metabolic syndrome (MetS) is a chronic condition that affects more than 1 billion people globally and a strong risk factor for knee osteoarthritis (OA) incidence, progression, and pain (*1–6*). Abnormal biomechanical stress on joint tissues due to obesity may partially mediate this association, yet MetS also increases the risk of hand OA, suggesting it also drives OA risk through systemic metabolic effects acting on joint tissues (*7*). As organs, synovial joints are designed to withstand biomechanical and metabolic stresses, but chronic exposure to stresses that exceed physiologically tolerated ranges contributes to failure of the joint organ system. This results in pain and loss of function for patients suffering from OA (*8*). Identifying the mechanisms driving these effects is a key objective to enable the development of disease-modifying OA treatments (*9*).

The synovial joint lining (synovium) plays a crucial role in maintaining joint organ health by providing nutrients (oxygen and macromolecules) to the avascular cartilage, joint lubrication, and immune surveillance. Given its pivotal role in synovial joint organ homeostasis and the association of synovial inflammation and damage with knee OA pain, we postulate that synovial cell or tissue dysfunction may lead to worse OA outcomes (*10–12*). Since metabolic stress drives dysfunction in other organs such as the kidney, we hypothesize that metabolic stress may also contribute to synovial tissue dysfunction. However, the separate and combined effects of biomechanical and metabolic stresses on synovial cells and tissues are not well understood.

The synovium is largely comprised of specialized macrophages and fibroblasts. These cells collaborate in the maintenance of tissue homeostasis through coordinated intercellular communication and mediate pathophysiological changes in OA (*13*). For example, the OA synovial fibroblast secretome has been implicated in pain and plays an important role in joint tissue damage (*14–16*). In addition, alteration of synovial macrophage activation during OA progression can alleviate pain and slow disease progression, whereas their ablation leads to worse inflammation and synovial tissue damage (*17, 18*). This highlights the importance of macrophages to OA outcomes and joint homeostasis. Early evidence suggests that metabolic stress worsens synovial inflammation and shifts macrophage activation toward a proinflammatory state (*19, 20*). Despite this, it still remains unclear how metabolic stress affects synovial cells and impacts outcomes in OA. Therefore, our objective was to investigate the separate and combined effects of biomechanical and metabolic stresses in OA, with a focus on changes occurring in synovial macrophages and fibroblasts.

In a cohort of patients with early-stage knee OA, we investigated the association between MetS and worse patient-reported knee pain while adjusting for potential confounders. This led us to test the effects of metabolic and/or biomechanical stresses in a rat model of experimental knee OA (*21, 22*). Herein, we demonstrate complementary and synergistic effects of biomechanical (induced by surgical joint destabilization) and metabolic (induced by high-fat and high-sucrose diet) stresses on pain, cartilage damage, and synovial histopathology. Further, we used single cell RNA sequencing (scRNA-seq) to investigate these mechanisms in synovial macrophages and fibroblasts, which likely play key effector roles on OA outcomes.

## Results

### Association of metabolic syndrome with clinical outcomes

A total of 153 knees from 153 patients were included. Two patients did not have pain scores available. Demographics and clinical characteristics are shown in Figure 1A.

**Figure 1.**
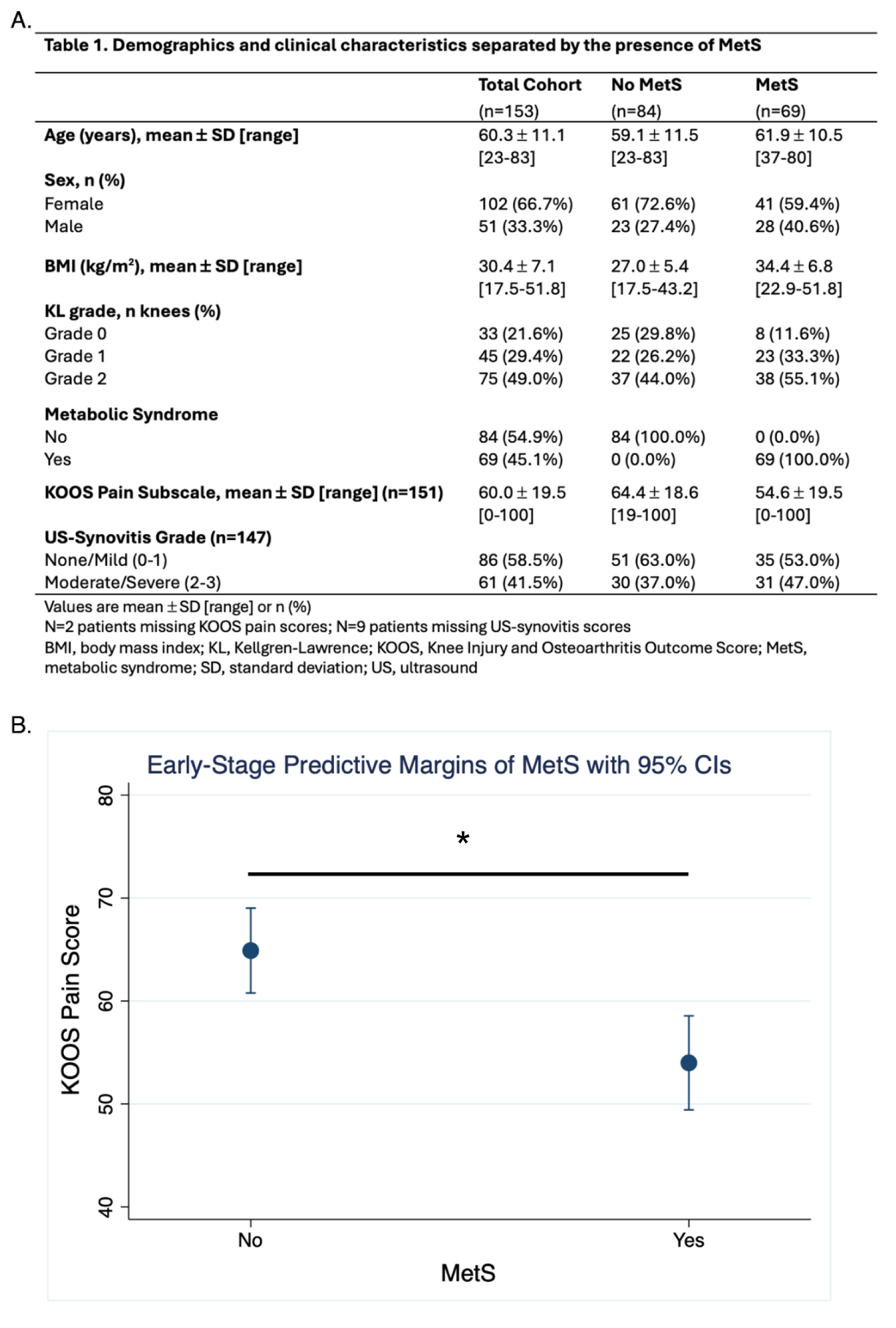
Clinical association of MetS with pain in early stage knee OA. Patient demographics (**A**) and clinical associations of KOOS pain with MetS in early stage knee OA (**B**). Y-axis represents KOOS pain and x-axis the presence or absence of MetS. Data are presented as unstandardized beta-coefficient ± 95% CI. * p< 0.05.

Metabolic syndrome was associated with lower Knee injury and Osteoarthritis Outcome Score (KOOS) pain scores (−10.90 [95%CI −17.11, −4.70]), indicating worse pain (Figure 1B; Supplementary Table 1) with an effect size that was likely clinically meaningful. Further, when synovitis was included in the model, the relationship between MetS and KOOS pain remained statistically significant (−9.70 [95%CI −16.15, −3.26] (Supplementary Table 1), indicating that MetS is an independent predictor of pain.

### Metabolic stress worsens pain, cartilage degeneration, and synovial inflammation in experimental knee OA

Based on these findings, we next sought to investigate a potential causal relationship between metabolic syndrome and pain in early-stage OA. We used a rat model of early-stage experimental knee OA to investigate the effects of metabolic stress on synovial tissues and OA outcomes. Conventionally, rodents are fed a lean diet (regular diet, RD) and OA is induced with biomechanical stress alone through surgical destabilization of the knee joint. Here, we compared healthy or biomechanically stressed animals to animals that underwent induction of metabolic stress through 12-weeks of feeding with a high-fat, high-sucrose diet (HFS), followed by the usual surgical induction of biomechanical stress. This allowed us to assess the effects of biomechanical stress, metabolic stress, and the combination during the development of experimental knee OA.

As expected based on our prior research with this model, induction of biomechanical stress through joint destabilization resulted in a transient reduction in knee withdrawal threshold (worse pain) at 4-weeks, which recovered to baseline at 8- and 12-weeks post-surgery. Interestingly, prior to the induction of biomechanical stress (baseline), the metabolic stress group (HFS) had a higher knee withdrawal threshold when compared to the regular diet group (RD) (127.67g [95%CI 32.52, 222.82], p<0.05) (Figure 2A-B). At the earliest time-point (4-weeks), the combination of metabolic and biomechanical stress exacerbated the magnitude of reduction in knee withdrawal threshold compared with biomechanical stress alone, consistent with worse pain and similar to our clinical findings (−243.26 grams [−297.36, −189.17] vs −68.73 grams [−130.52, −6.94] respectively) (Figure 2A and Supplementary table 2). In contrast to biomechanical stress alone, animals with combined metabolic and biomechanical stress had persistent mechanical pain sensitivity through 8- (−121.05 grams [−176.04, −65.78]) and 12-weeks (−157.95 grams [−212.94, −102.68]). Both groups experienced a reduction in ipsilateral hindpaw withdrawal threshold at 4-weeks, that was maintained through the 8- and 12-week timepoints (Figure 2C-D and Supplementary table 3). Taken together, these data suggest that metabolic stress may exacerbate and prevent the resolution of mechanical knee pain sensitivity in early-stage knee OA.

**Figure 2.**
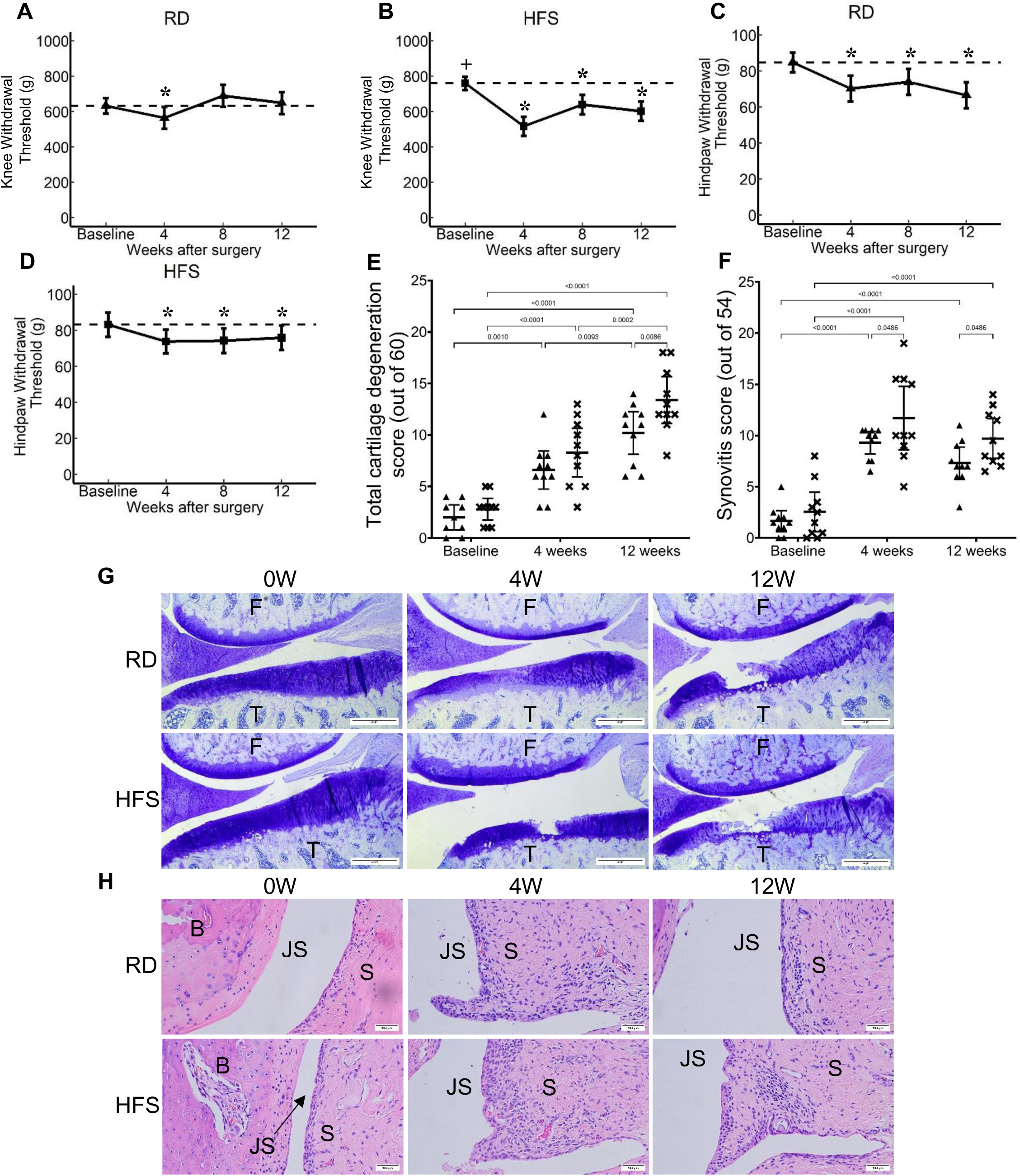
Characterisation of experimental rodent model. Pressure pain threshold of RD (**A**) or HFS (**B**) and hindpaw withdrawal threshold of RD (**C**) or HFS (**D**) at baseline, and, 4-, 8-, and 12-weeks post OA induction. Y-axis represents force (grams, g) applied to the knee or ipsilateral hindpaw before withdrawal and x-axis time in weeks. Mean with 95% CI are displayed. Significant differences versus baseline (*) and versus RD (+) are shown (p<0.05). Measures of cartilage degeneration in the whole joint of RD (▲) or HFS (**x**) at baseline, 4-, and 12-week timepoints (**E**). Y-axis shows the total histopathological score (out of 60) and x-axis time in weeks. Mean with 95% CI are displayed. Measures of synovial inflammation in the whole joint of RD (▲) or HFS (**x**) at baseline, 4-, and 12-week timepoints (**F**). Y-axis shows the total histopathological score (out of 54) and x-axis time in weeks. Mean with 95% CI are displayed. Representative histological images of toluidine blue stained medial cartilage (**G**). F denotes femur and T tibia. Scale bar = 500µm. Representative H&E stained synovial histological images (**H**). B denotes bone, JS joint space, and S synovium. Scale bar = 100µm.

Cartilage degeneration was similar between groups prior to induction of biomechanical stress (Figure 2E). Following joint-destabilization, cartilage degeneration progressed in severity at 4- and 12-weeks and was marked by proteoglycan loss, fissuring, and partial thickness erosions, consistent with previous literature. The combination of metabolic and biomechanical stress led to worse cartilage degeneration at 12-weeks when compared to biomechanical stress alone.

Synovial inflammation (synovitis) was similar between groups prior to joint destabilization and worsened in both groups at 4- and 12-weeks (Figure 2F). Interestingly, the combination of metabolic and biomechanical stress increased the severity of synovitis at both 4- and 12-weeks when compared to biomechanical stress alone (p<0.05) with the differences at 12-weeks being driven primarily by lining hyperplasia and fibrosis (Supplementary Figure 1). Taken together with our clinical findings, these results confirm that metabolic stress exacerbates pain and structural joint damage in the early stages of experimental knee OA.

### Single-cell transcriptomics profiling of synovium reveals metabolic stress associated cellular stress and oxidative damage

We next used scRNA-seq to define and investigate the effects of biomechanical, metabolic, and combined biomechanical and metabolic stress on synovial cell populations. Using previously described gene markers, we found 21 cell clusters present in all groups (Figure 3A and B, Supplementary figure 2). As expected, we found all major cell types including fibroblasts, myeloid cells, endothelial cells, pericytes, T cells, B cells, and Schwann cells (*16*). In healthy synovium (RD 0W), fibroblast and myeloid cells comprised the largest proportion of cells (Figure 3C). The proportion of fibroblasts increased and myeloid cells decreased in response to biomechanical stress alone (4-week time-point, RD 4W), and both cell types recovered to near-baseline levels by 12-weeks (RD 12W). At baseline, metabolic stress (HFS 0W) slightly increased the proportion of fibroblasts when compared to regular diet (RD 0W). When metabolic and biomechanical stress were combined, fibroblasts increased at 4- (HFS 4W) and 12-weeks (HFS 12W), while myeloid cells decreased at 4-weeks before returning to baseline. These data suggest that control of fibroblast expansion may be impaired by metabolic stress contemporarily with the onset of persistent mechanical pain sensitivity.

**Figure 3.**
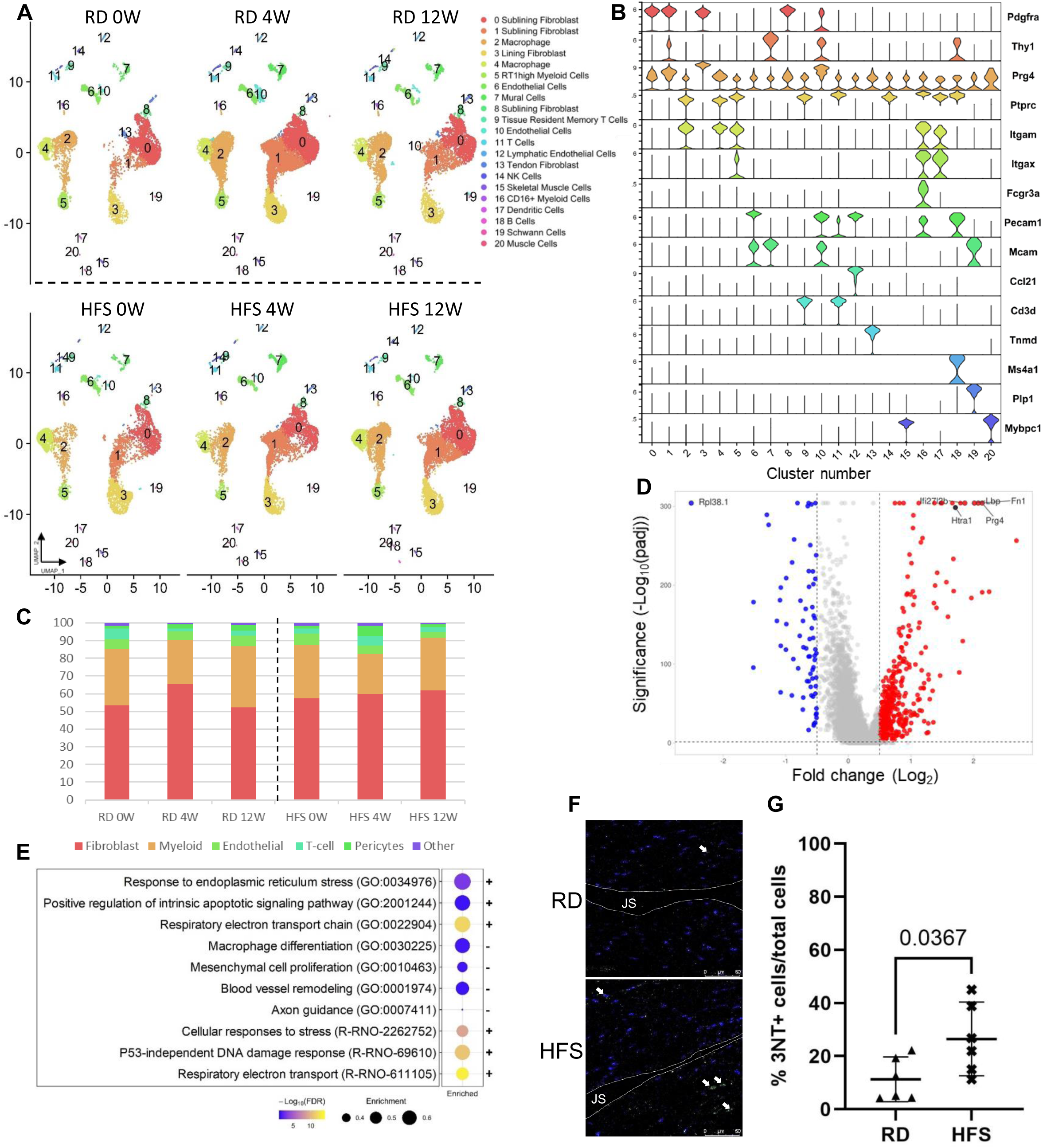
Single cell transcriptomic profiling of synovium. UMAP plots showing projections of synovial cells from RD or HFS at baseline (0W), 4- (4W), or 12-weeks (12W) (**A**). Cells types are grouped by color with groups designated by number and annotations on the right. Violin plots of synovial cell marker genes (**B**). Proportional breakdown of major synovial cell types in each condition (**C**). Volcano plot showing the differentially expressed genes between synovial cell at RD 0W and HFS 0W (**D**). Y-axis represents the −log10 of the adjusted p-value with a cut-off of 1.3 (padj <0.05) and x-axis the log2 fold change with cut-off at −0.5 and 0.5. Bubble plot showing curated gene set enrichment of HFS 0W compared to RD 0W (**E**). Enrichment represents the number of genes called to a gene set divided by the total number of genes in the set (**F**). Abundance of 3NT+ cells in the synovium of RD (▲) or HFS (**x**) at baseline (**G**). Y-axis represents 3NT+ cells as a percentage of total cells and x-axis diet.

To investigate the effects of metabolic stress alone on the transcriptional profile of all synovial cells, we used pseudobulk analysis to identify differentially expressed genes (DEGs) and performed pathway analyses. At baseline, we identified 487 DEGs enriched in metabolic stress when compared to healthy controls, with 70 reduced and 417 increased (Figure 3D, Supplementary table 4). Top dysregulated synovial cell genes in metabolic stress included Htra serine protease 1 (Htra1), fibronectin 1, lubricin, lipopolysaccharide binding protein, and 60S ribosomal protein L38. Next, pathway enrichment identified p53-dependent DNA damage, endoplasmic reticulum stress, cellular stress, apoptosis, and energy production pathways were increased by metabolic stress (Figure 3E and Supplementary table 5-6). Further, macrophage differentiation, mesenchymal cell proliferation, blood vessel remodeling, and axon guidance pathways were all decreased in metabolic stress.

Since cell and endoplasmic reticulum stress pathways can be triggered by or lead to oxidative stress, we next assessed the abundance of cells positive for 3-nitrotyrosine (3NT+) to confirm that metabolic stress increased oxidative stress in synovial cells (*23, 24*). Indeed, 3NT+ synovial cells were increased at baseline in metabolic stress when compared to healthy (24.3% positive cells [95%CI 13.4-35.2] vs 11.3 [2.5-20.2] respectively, p=0.0392) (Figure 3F-G). These transcriptomic and in situ data suggest that metabolic stress induces oxidative stress in synovial cells, which may underlie aberrant synovial cell activation and dysfunction.

### Metabolic stress is associated with transcriptional programs of fibroblast activation

Fibroblasts are the most abundant synovial cell type and are associated with worse pain in knee OA (*14*). To understand the effect of metabolic stress on synovial fibroblasts, we computationally subset and re-clustered them, identifying 7 subsets including Lrrc15+, Pi16+/Dpp4+ progenitors, Prg4-high lining (herein referred to as lining), Dpp4+ progenitor, Il6+/Ccl2+, perivascular, and tendon fibroblasts (Figure 4A and Supplementary Figure 3A-C).

**Figure 4.**
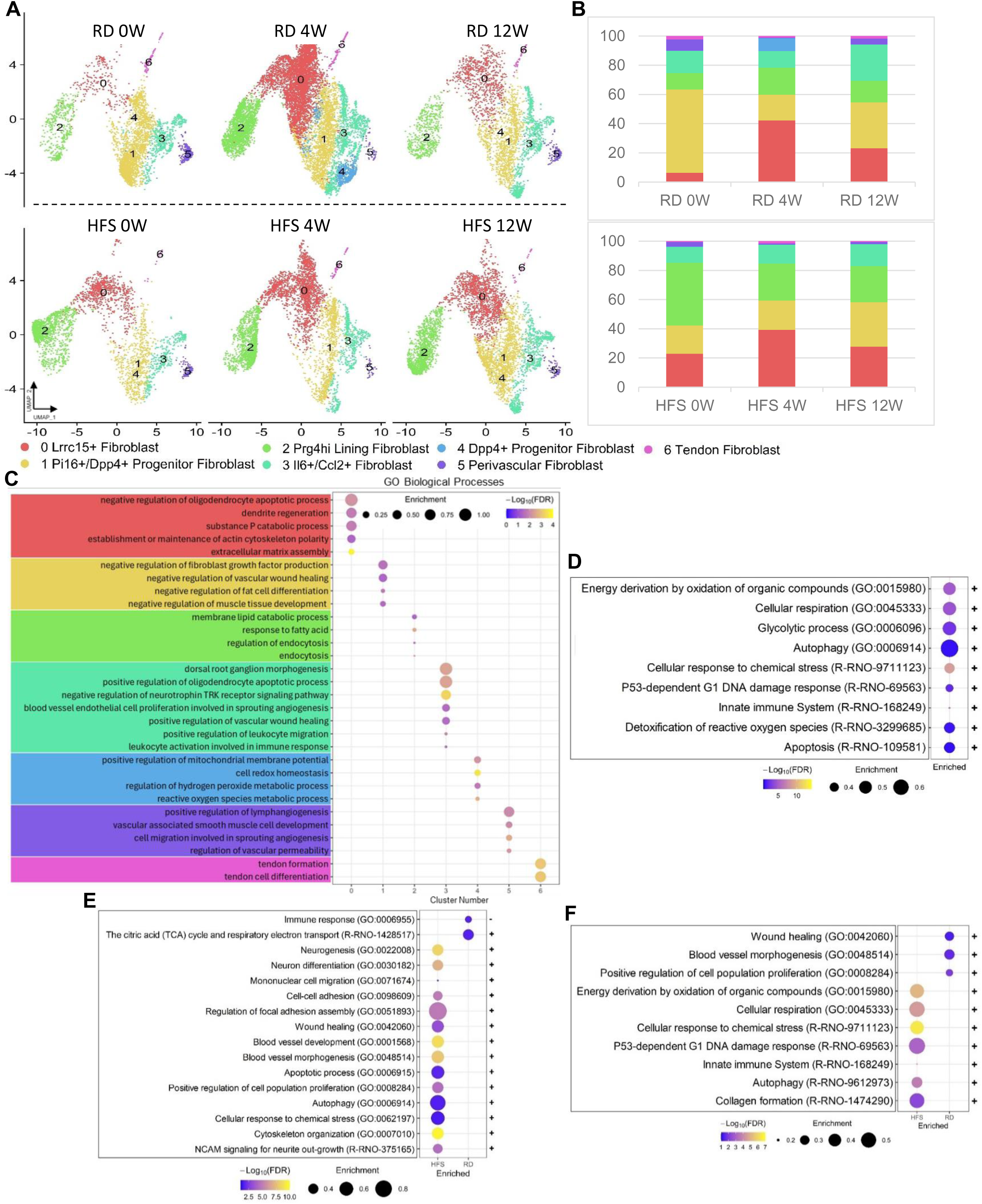
Single cell transcriptional profiling of synovial fibroblasts. UMAP plots showing projections of synovial fibroblasts from RD or HFS at baseline (0W), 4- (4W), or 12-weeks (12W) (**A**). Fibroblast subsets are grouped by color with groups designated by number and annotations on the right. Proportional breakdown of fibroblast subsets in each condition (**B**). Pathway analysis using Gene Ontology (GO) Biological Processes showing unique terms of each cluster (**C**). Pathway analysis using Gene Ontology (GO) Biological Processes and Reactome comparing baseline HFs to RD (**D**). Pathway analysis using Gene Ontology (GO) Biological Processes and Reactome showing unique terms of RD or HFS at 4- and 12-week OA timepoints (**E-F**).

Pi16+/Dpp4+ progenitors were the largest subset of synovial fibroblasts in healthy synovium, but decreased in response to biomechanical stress, metabolic stress, and the combination, and only recovered partially by 12-weeks (Figure 4B). Functional annotations identified pathways associated with negative regulation of growth factor signaling, wound healing, cell differentiation, and tissue development, suggesting a potential role in wound healing resolution and maintenance of homeostasis (Figure 4C).

Lrrc15+ fibroblasts were sparse in healthy synovium but expanded in response to biomechanical stress, metabolic stress, and the combination and remained abundant at 12-weeks. Functional annotations identified pathways associated with regulation of dendrites and associated oligodendrocytes and extracellular matrix assembly, suggesting functional roles in neurogenesis.

Lining fibroblasts were the third largest subset in healthy synovium and expanded in response to biomechanical stress before returning close to healthy levels at 12-weeks. Surprisingly, lining fibroblasts dramatically expanded in response to metabolic stress (43% metabolic stress vs. 11% healthy), but contracted when metabolic and biomechanical stress were combined (43% metabolic stress vs. 25% metabolic plus biomechanical stress). Functional annotations identified pathways associated with endocytosis, suggesting these cells may promote joint homeostasis through removal of joint products.

Il6+/Ccl2+ fibroblasts were the second largest subset in healthy synovium, but contracted in response to the combination of biomechanical and metabolic stress. Interestingly, by 12-weeks, Il6+/Ccl2+ fibroblasts expanded in response to biomechanical stress alone, corresponding to the resolution of mechanical knee pain sensitivity, milder synovitis, and less cartilage degeneration. Functional enrichment identified pathways involved in immune signaling and activation, wound healing, and pain processes, suggesting a potential pro-regenerative role.

Dpp4+ progenitors were primarily present at 4-weeks in response to biomechanical stress alone with functional annotations associated with redox processes suggesting these cells may aid in resistance to oxidative stress. Perivascular fibroblasts diminished in response to biomechanical stress, metabolic stress, and the combination, and were functionally associated with maintenance and regulation of blood and lymphatic vessels. Lastly, tendon fibroblasts were primarily associated with tendon formation pathways.

Pathway enrichment analysis in fibroblasts demonstrated that metabolic stress increased pathways associated with energy production, cell stress, apoptosis, and DNA damage (Figure 4D, Supplementary table 7-9). Next, we interrogated pathways unique to either biomechanical or combined metabolic and biomechanical stress. At 4-weeks, pathways associated with reduced immune signaling and increased energy production were enriched by biomechanical stress, whereas pathways associated with neuronal development, leukocyte recruitment, wound healing, blood vessel formation, apoptosis, and autophagy were enriched when metabolic and biomechanical stress were combined (Figure 4E, Supplementary table 10-15). At 12-weeks, pathways associated with wound healing and blood vessel formation were enriched by biomechanical stress alone, whereas pathways associated with energy production, cellular stress, and collagen formation were enriched when metabolic and biomechanical stress were combined (Figure 4F, Supplementary tables 16-21). Taken together with the differential effects on synovial cell subsets, these data suggest that metabolic stress shifts synovial fibroblasts away from negative regulation of cell activation, and induces endocytic, vascular remodeling, and neurogenic functions.

### Metabolic stress increases myeloid cell stress and causes metabolic shifts

Re-clustering of myeloid cells identified 11 populations including Cd163+ interstitial macrophages, synovial lining macrophages, MHCII high dendritic cells, MHCII high macrophages, Aqp1+ interstitial macrophages, monocytes, plasmacytoid dendritic cells, conventional type 1 dendritic cells, mature dendritic cells, proliferative myeloid cells, and mast cells (Figure 5A, Supplementary Figure 4A-C). Cd163+ interstitial macrophages were the most abundant myeloid cell in healthy synovium, which reduced in response to biomechanical stress before returning to similar levels at 12-weeks (Figure 5B). Surprisingly, metabolic stress reduced the abundance of Cd163+ macrophages substantially (9% Metabolic stress vs 44% healthy) but when biomechanical stress was combined they expanded to proportions similar to animals without metabolic stress. Functional annotations identified pathways associated with synapse pruning and regulation of neuronal support cells, negative regulation of vascular smooth muscle cells, positive regulation of endothelial cells, and negative regulation of nitric oxide biosynthetic processes and type II interferon. These data suggest that Cd163+ synovial macrophages play roles in wound healing and limiting tissue damage (Figure 5C).

**Figure 5.**
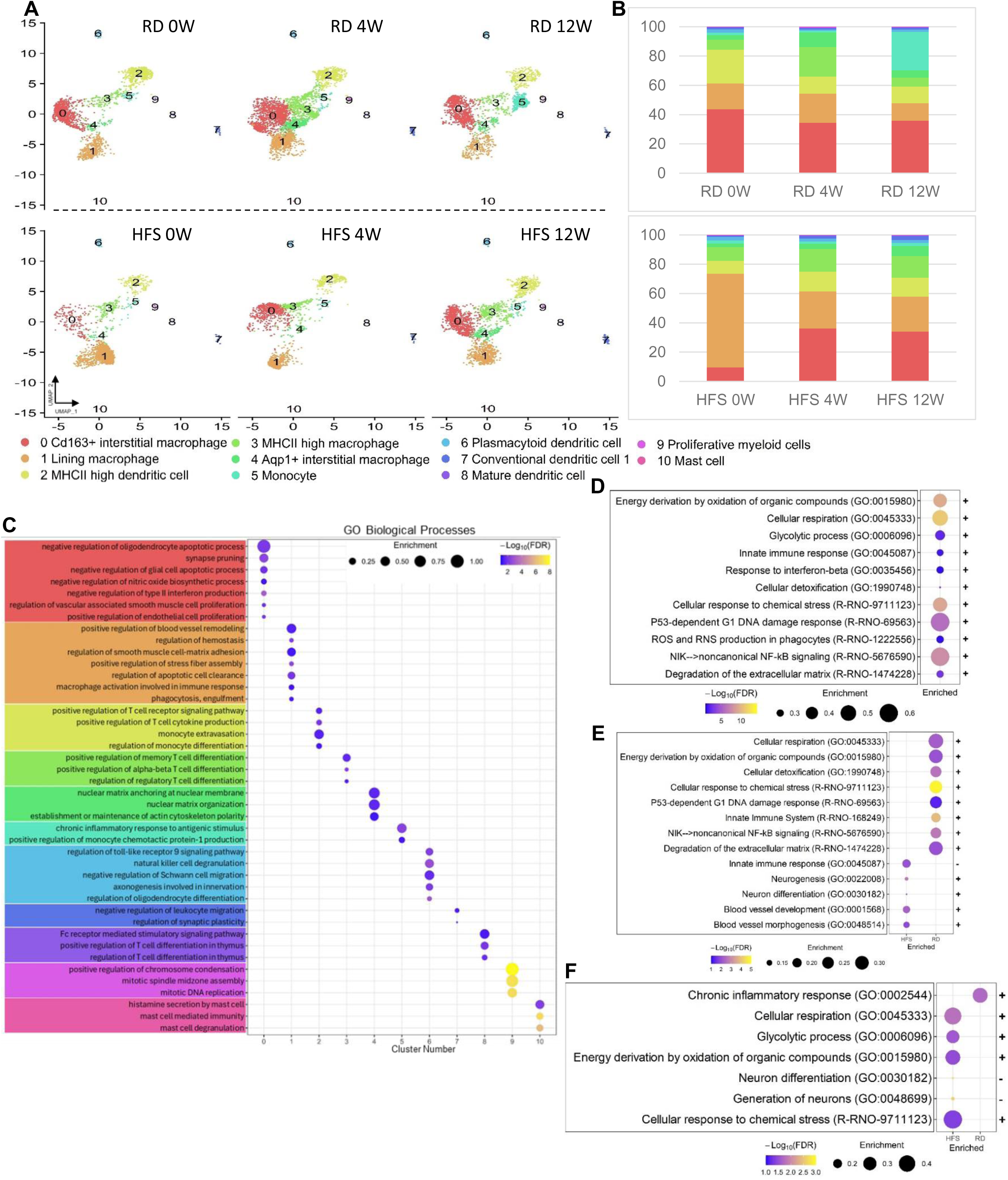
Single cell transcriptional profiling of synovial myeloid cells. UMAP plots showing projections of synovial myeloid cells from RD or HFS at baseline (0W), 4- (4W), or 12-weeks (12W) (**A**). Myeloid subsets are grouped by color with groups designated by number and annotations on the right. Proportional breakdown of myeloid subsets in each condition (**B**). Pathway analysis using Gene Ontology (GO) Biological Processes showing unique terms of each cluster (**C**). Pathway analysis using Gene Ontology (GO) Biological Processes and Reactome comparing baseline HFs to RD (**D**). Pathway analysis using Gene Ontology (GO) Biological Processes and Reactome showing unique terms of RD or HFS at 4- and 12-week OA timepoints (**E-F**).

Lining macrophages were the third largest population of myeloid cells in healthy synovium. Their proportion increased 4-weeks after induction of mechanical stress but reduced below normal levels by 12-weeks. After induction of metabolic stress, lining macrophages dramatically expanded (64% metabolic stress vs 18% healthy), but contracted when biomechanical stress was added, corresponding to worse outcomes. Functional annotations identified pathways related to homeostasis, phagocytosis, efferocytosis, and regulation of vessel remodeling, suggesting that lining macrophages may play a role in maintaining tissue homeostasis.

MHCII high dendritic cells were the second most abundant myeloid cell in healthy tissue, but were reduced in response to biomechanical stress, metabolic stress, and the combination. Functional annotations identified pathways associated with T-cell activation and cytokine production, suggesting these cells play a role in regulating T-cell function in the joint.

Aqp1+ macrophages increased in abundance in response to biomechanical stress but returned to baseline levels at 12-weeks. Interestingly, the expansion of Aqp1+ macrophages was delayed to 12-weeks when metabolic stress was combined with biomechanical stress. Functional annotations identified pathways related to nuclear matrix organization and cytoskeletal organization, suggesting these cells may be migratory and adjusting shape and polarity.

Infiltrating monocytes expanded at 12-weeks in response to biomechanical stress alone, but this was blunted when metabolic and biomechanical stress were combined. Functional enrichment identified pathways associated with chronic response to antigenic stimulus and regulation of C-C motif chemokine ligand 2 production. Functional enrichment in plasmacytoid dendritic cells identified pathways related to axonogenesis and differentiation and migration of axon supporting cells, suggesting potential roles in joint nociception. Conventional type 1 dendritic cell functional annotations identified pathways related to negative regulation of leukocyte migration, while mature dendritic cells were associated with pathways associated with T-cell differentiation. Lastly, proliferative myeloid cells were associated with DNA replication pathways and mast with degranulation and histamine secretion.

Enrichment analysis identified pathways related to glycolysis and respiration, detoxification, oxidative stress, cell stress, DNA damage, innate immune responses, and reactive oxygen and nitrogen generation when compared to healthy synovial myeloid cells (Figure 5D and Supplementary tables 22-24). At 4-weeks, metabolic stress combined with biomechanical stress enriched pathways related to blood vessel formation and neurogenesis (Figure 5E and Supplementary tables 25-27). Biomechanical stress alone induced pathways related to metabolic shifts, detoxification, oxidative stress, cell stress, and innate immune response similar to those induced by metabolic stress alone (Supplementary tables 28-30). By 12-weeks, biomechanical stress alone enriched one unique pathway associated with chronic inflammatory response whereas, metabolic stress combined with biomechanical stress enriched pathways suggesting ongoing metabolic shifts, cell stress, and neurogenesis (Figure 5F and Supplementary tables 31-36). These data suggest that metabolic stress drives shifts in cellular metabolism and increases cellular stress, while prolonging the activation of neurogenesis.

### Macrophage to fibroblast communication is disrupted by metabolic stress

Cellular communication, especially between macrophages and fibroblasts, is a critical component of homeostasis and wound healing. With this in mind, we first assessed all outgoing communication patterns in healthy and metabolically stress synovium. Four outgoing communication patterns emerged in healthy synovium (Figure 6A). One pattern was associated with fibroblasts, one with immune cells, and two with endothelial and vascular support cells. Unlike healthy synovium, metabolic stress increased the number of predicted outgoing communication patterns, with separate patterns for myeloid and lymphoid cells (Figure 6B). Interrogation of outgoing communication pathways identified 7 that were unique to metabolic stress, including SELL, NGL, SPP1, VISTA, NRG, Opioid, and XCR. Myeloid cells contained the highest number of dysregulated outgoing pathways (4 of 7), suggesting that metabolic stress may readily alter myeloid communication.

**Figure 6.**
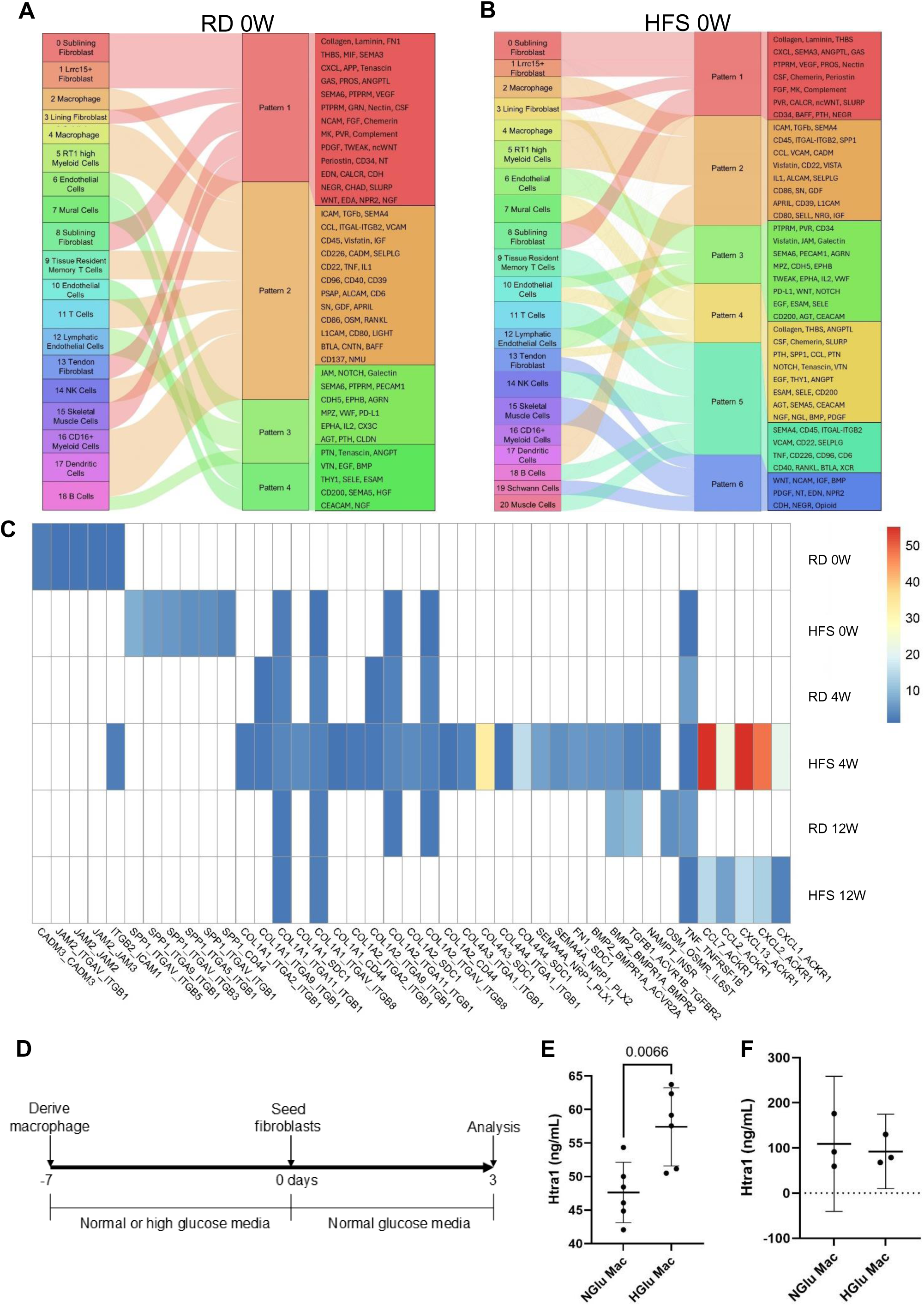
Profiling of cellular communication. River plots showing outgoing communication patters from all synovial cells at baseline in RD (**A**) or HFS (**B**) conditions. Heatmap of altered macrophage to fibroblast ligand receptor interactions (**C**). Color scale shows probability ratio. Schematic of metabolic stress co-culture experiment (**D**). Htra1 production by macrophage-fibroblast culture system (**E**). NGlu Mac denotes cultures containing macrophages derived in normal glucose media and HGlu Mac cultures containing macrophages derived in high glucose media. Htra1 production by normal (NGlu Mac) or high (HGlu Mac) glucose derived macrophages (**F**).

Considering this and the importance of macrophage-to-fibroblast communication, we next explored macrophage to fibroblast ligand-receptor (LR) interactions that were unique to healthy synovium or enriched in stressful conditions. Ligand receptor interactions associated with cell adhesion, junctional adhesion, and intracellular adhesion molecules were unique to healthy synovium with the exception of Itgb2-mediated signaling, which was found at 4-weeks when metabolic stress was combined with biomechanical stress (Figure 4C). Biomechanical stress induced matrisomal (collagen 1 [COL1]) and inflammatory (tumor necrosis factor [TNF]) LR interactions at 4-weeks and inflammatory (Oncostatin M and TNF), matrisomal (COL1), and transforming growth factor beta superfamily (bone morphometric protein 2 [BMP2] and transforming growth factor beta-1 [TGFβ1]) LR interactions at 12-weeks. Metabolic stress alone induced Spp1 mediated LR interactions. Combining metabolic and biomechanical stress induced matrisomal (COL1 and 4, and fibronectin 1), semaphorin, TGFβ superfamily (BMP2 and TGFβ1), adipokine (Visfatin), inflammatory (TNF), and chemokine LR interactions. At 12-weeks, matrisomal (COL1) and chemokine LR interactions remained enriched in the combined group. Interestingly, when metabolic and biomechanical stress were combined, communication networks that promote activation of fibroblasts and fibrosis (TGFβ1, Visfatin, and semaphorin) were induced sooner then in biomechanical stress alone.

Given the enrichment of adhesion-mediated cellular communication in healthy synovium, cell-cell contact between macrophages and synovial fibroblasts may be important for controlling synovial fibroblast behaviour. Therefore, we next used a direct contact macrophage-fibroblast co-culture system to explore whether metabolic stress induced changes in macrophage-fibroblast communication alters the production of Htra1 a top DEG induced by metabolic stress and contributor to OA progression (*25*). To induce metabolic stress, blood-derived macrophages were cultured in high glucose media (HGlu Mac) for 7 days prior to co-culture with synovial fibroblasts in normal glucose media (Figure 6D). When OA synovial fibroblasts were cultured in direct contact with HGlu Macs, they produced significantly more Htra1 than fibroblasts co-cultured with healthy macrophages (NGlu Mac) (50.5ng/mL [95%CI 51.6-63.3] and 42.1ng/mL [43.1-52.1], respectively) (Figure 6E). We confirmed that metabolic stress did not alter the production of Htra1 by macrophages (Figure 6F). These data suggest that exposure of macrophages to metabolic stress alters macrophage-fibroblast communication, leading to increased Htra1 production.

## Discussion

Metabolic syndrome is a strong risk factor for OA onset and progression. The systemic effects of metabolic stress on synovial cell populations, either alone or in combination with biomechanical stresses, may lead to worse OA outcomes through mechanisms that need to be better understood. Here, we discovered that MetS (a source of metabolic stress on joint tissues) is independently associated with worse knee pain in early-stage knee OA. This suggests that MetS may exacerbate pain and OA progression, but the underlying mechanisms are unclear. We then used a rat model of experimental knee OA to compare and contrast the effects of biomechanical and metabolic stresses on joint outcomes. When combined with biomechanical stress, metabolic stress induced worse and sustained knee pain, cartilage damage, and synovial tissue inflammation and damage, supporting the hypothesis that metabolic stress plays a causal role in worsening OA outcomes.

Using single-cell transcriptional profiling, we found that metabolic stress and biomechanical stress both independently activated glycolytic, respiratory, and oxidative metabolic pathways in synovial macrophages, increased the abundance of homeostatic lining cells, induced wound healing pathways in synovial fibroblasts, and disrupted adhesion-mediated macrophage-to-fibroblast communication. In isolation, metabolic stress increased oxidative stress, and activated glycolytic, respiratory, and oxidative metabolic pathways in fibroblasts. Combining metabolic and mechanical stress induced neurogenic pathways in both fibroblasts and macrophages. Further, the combination induced macrophage-to-fibroblast communication through adipokine, chemokine, and semaphorin interactions. Interestingly, the combination also induced metabolic alterations, wound healing, and TGFβ superfamily communication approximately 8 weeks earlier when compared to biomechanical stress alone. Lastly, in vitro co-cultures showed altered communication from metabolically stressed macrophages to synovial fibroblasts led to increased production of Htra1, a suspected driver of OA. Taken together, these data suggest that early dysregulation of synovial cells, their communication, metabolism, and wound healing processes may be key events during early-stage OA that can lead to worse OA outcomes.

Preclinical models of metabolic stress have been developed employing dietary formulations of varying fat and carbohydrate concentrations. Similar to others, we found that pain, cartilage damage, and synovitis were worsened when metabolic and mechanical stress were combined (*18, 20, 26*). In line with Wu et al., but unlike Collins et al. and Sun et al., we did not identify a difference in cartilage degeneration or synovitis in metabolic stress alone (*19, 21, 26*). However, we did see trends towards worse cartilage and synovial pathology prior to OA induction. Differences in dietary formulations, timepoints, histopathological scoring systems, animal age, and animal strains may underlie variation between studies. Nonetheless, our histopathological findings are generally aligned with similar studies in the field.

Osteoarthritis pain is a multidimensional construct that is strongly associated with radiographic severity (cartilage and bone damage), but factors aside from structural damage also have important influences on pain outcomes. For example, synovial inflammation (synovitis), which develops prior to radiographic bone and cartilage damage, is also associated with increased severity of pain in OA (*10, 11*). Further, we and others have shown that macrophages and fibroblasts contribute to nociception in OA, strongly suggesting the synovium plays a critical role in OA pain (*17, 27*). In our cohort study, we found that MetS is associated with worse pain in early-stage knee OA where there is minimal to no radiographic damage. Although the association was partially confounded by synovitis, it nonetheless remained present with a large enough effect size to be clinically-meaningful after adjusting for synovial inflammation. This suggested to us that metabolic stress may also be an important contributor to worse pain experiences in early-stage knee OA independent from other well-known factors. Similarly, the combination of metabolic and biomechanical stresses in our preclinical model of experimental OA led to worse pain that was sustained. Further, transcriptional profiling of synovial fibroblasts and macrophages identified enrichment of pathways associated with neurogenesis, neuron differentiation, and generation of neurons. Supporting this, fibroblasts from both OA and rheumatoid arthritis patients have been shown to promote neurogenesis in vitro (*14, 28*). Together, these data suggest synovial fibroblasts and macrophages may mediate the effects of metabolic stress leading to worse pain experiences during the early stages of OA development and progression.

Our transcriptomic data suggested that metabolic stress increased the abundance of synovial cells experiencing oxidative stress, which we confirmed with immunofluorescence detection of 3NT+ cells in synovial tissue. Cellular damage from excess oxidative stress is a central player in the induction of apoptosis (*29*). Interestingly, pathway enrichment identified increased synovial fibroblast apoptosis after induction of metabolic stress. If apoptotic cells are not removed efficiently by phagocytes via efferocytosis, they can contribute to pain and inflammation through release of damage associated molecular patterns. We have previously shown that efferocytosis by synovial macrophages is impaired in OA and may contribute to worse outcomes (*30*). In our model, lining macrophages demonstrated functional terms including homeostasis and efferocytosis. These cells became more abundant in response to metabolic stress, but reduced when biomechanical stress was added, corresponding to worse pain and synovitis. Taken together, these data suggest that lining macrophages may respond acutely to stresses but can be lost when exposed to chronic stresses, which might be an important factor determining OA outcomes.

The extracellular matrix (ECM) is a major regulator of cell and tissue function (*31*). Both fibroblasts and macrophages play vital roles in the maintenance of the ECM through tissue remodeling, which is coordinated by intercellular communication and involves the breakdown and deposition of ECM molecules (*13*). Disrupted tissue remodeling mechanisms can lead to synovial fibrosis (excess ECM), which contributes to reduced joint function in OA (*32*). We found that metabolic and mechanical stress synergistically worsened synovial fibrosis at 12-weeks, which overlapped with transcriptional enrichment of the collagen formation pathway. Interestingly, this occurred after enrichment of macrophage-to-fibroblast ligand receptor interactions that promote fibroblast activation and fibrosis (TGFβ1, Visfatin, and semaphorin) (*33–35*). Fibroblast activation due to dysregulated macrophage-to-fibroblast communication may have contributed to the synovial fibrosis identified here.

We found that joint stress (metabolic, biomechanical, or both) reduced the abundance of cell-cell adhesion-mediated communication. Although little is known about adhesion molecule-mediated communication in the context of macrophages and fibroblasts, engagement of tight junctions regulates proliferation, cellular activation, and maintenance of barriers (*36–38*). Interestingly, Culemann and colleagues found that ablation of tight junction-expressing synovial lining macrophages led to disruption of the lining architecture (*39*). In addition, we recently identified erosion of the synovial lining in late-stage knee OA tissue and found that it associates with other features of synovial damage (*12*). It is possible that loss of adhesion-mediated communication may represent an early step in synovial lining erosion that warrants future exploration.

Catabolic breakdown of joint tissue during tissue remodeling is facilitated by proteases including Htra1, a serine peptidase that cleaves aggrecan, fibronectin, and biglycan. Htra1 increases in OA, where its proteolytic function can worsen outcomes (*25*). Fibronectin fragments produced by Htra1 promote inflammation, a known contributor to OA pain, and cause release of metalloproteinases that drive cartilage damage (*40, 41*). Further, Htra1 facilitates secretion of type 1 collagen and promotes activation of fibroblasts, leading to fibrosis (*42, 43*). Our pseudobulk analysis identified Htra1 as a top differentially increased gene in metabolic stress. At the same time, we found that metabolic stress reduced intercellular junctional adhesion-mediated communication, which may regulate fibroblast activation. In addition, we found that metabolic stress dysregulates macrophage-fibroblast communication and causes increased production of Htra1 using an in vitro assay that incorporated cell contact. Although these effects may have been due to contact-dependent or -independent communication (or both), our results suggest that maintaining cell adhesion-mediated communication may reduce Htra1 production.

Our study has limitations. We did not explore which molecular factors might be responsible for the effects of metabolic stress, although our in vitro high glucose assay suggests that hyperglycaemia may play a role and should be investigated further. Our surgically-induced model of OA did not address female sex, a risk factor for OA, and therefore we cannot rule out any effects of sex on the relationship between metabolic stress and experimental outcomes. However, our clinical cohort included a majority of female sex participants, suggesting that the association between MetS and pain is relevant to females and males. Our study also had strengths. Our preclinical model allowed us to investigate a causal relationship between metabolic stress and pain that was suggested by our findings of a clinical association in patients with early-stage knee OA. Further, we incorporated multiple OA risk factors including age and metabolic stress into our preclinical model, which may more accurately represent the complexity of the human disease, increasing the translatability of our findings.

In conclusion, MetS is independently associated with pain in early-stage knee OA. Further, in response to biomechanical stress, animals pre-exposed to metabolic stress had worse pain, and synovial inflammation, and cartilage damage, which may be related to dysregulated intercellular communication between synovial macrophages and fibroblasts, promoting accelerated cellular activation and Htra1 production. These novel findings suggest that targeting intercellular communication between synovial macrophages and fibroblasts to may provide therapeutic benefits to patients experiencing OA and MetS.

## Methods

### Study design

The purpose of our study was to examine the mechanisms of metabolic stress that influence outcomes in OA. We used human clinical data to assess associations between metabolic syndrome and pain in early stage knee OA. Next, animal models of metabolic stress, biomechanical stress, and the combination of metabolic and mechanical stress were employed in conjunction with behaviour analysis, histopathological analysis, scRNA-seq analysis, and immunofluorescence to identify dysregulated mechanisms and their relationship to OA outcomes. A direct contact culture system was used to investigate in vitro communication between OA fibroblasts and metabolically stressed macrophages. Sample sizes were guided by a power analysis and endpoints were determined from previous studies. Male Sprague-Dawley rats were used and purchased from Charles River Laboratories. Animals were randomly assigned to both diet and endpoint groups. Collection and use of human clinical data and biological samples was approved by Western University’s Research Ethics Board for Health Sciences Research Involving Human Subjects (HSREB #109255). Informed consent was obtained from all participants. All animal testing was approved by the Western University Animal Care and Use Committee under protocol AUP2021-087. Histopathological grading was performed by blinded investigators. All collected data were used for analysis. Detailed methods can be found in supplementary methods.

### Statistical Analysis

A cross-sectional multivariable linear regression model was fit to evaluate the association of metabolic syndrome (No/Yes) with patient-reported pain scores. All analyses were adjusted for age and sex. In secondary analyses, we evaluated the association of metabolic syndrome with pain while adjusting for age, sex, and ultrasound-synovitis. All analyses were completed using Stata/SE 15.1 (StataCorp, College Station, Texas, USA). For linear regression, we reported results as unstandardized beta (β) coefficients with 95% confidence intervals (CIs).

Mechanical sensitivity analyses were longitudinal and analyzed using linear mixed effects regression models for each diet (RD or HFS). Assumptions for linear mixed models were tested and likelihood ratios tests and Bayesian Information Criterion were used to evaluate model fit. Estimates of association are reported as unstandardized β coefficients ± 95% confidence intervals (95% CI) and standard error. Post-estimate pairwise comparisons of HFS to RD were completed with Tukey multiplicity correction. All statistical analyses of mechanical sensitivity data were performed using R (v4.3.3) with lme4 (v1.1-35.1), lmerTest (v3.1-3) and emmeans (v1.10.0) packages.

Single cell RNA-sequencing analysis was performed in R using the default statistical tests built into the Seurat and CellChat packages. All other analysis was performed in Prism Graphpad (v10.2.2) using two-way analysis of variance (ANOVA) with Tukey multiplicity adjustment or unpaired t-test.

## Supplementary material

Supplementary methods

Supplementary figures 1 to 4

Supplementary tables 1 to 36

## Acknowledgements

The authors would like to acknowledge all members of the WOREO Knee Study research team as well as the participants in the WOREO Knee Study. We are also grateful to Dr. Tristan Maerz, Dr. Alexander Knights, and Easton Farrell for their support and expertise while building our single cell RNA sequencing pipeline. We would also like to thank Christopher Wong for help in identifying myeloid cluster markers.

## Author contributions

Study conception and design; GB, JK, CTA. Acquisition of data; GB, JK, QG, HTP, HJ, DO, LD. Analysis and interpretation of data; GB, JK, QG, HTP, BF, LAW, CTA. Original draft writing; GB, JK, CTA. All authors were involved in revising the manuscript critically for important intellectual content, approved the final version submitted, and take responsibility for the integrity of data and accuracy of data analysis as per ICJME guidelines.

## Data availability

Single cell RNA sequencing data files were deposited in the Gene Expression Omnibus (GEO) at the NCBI (GSE273843). All other datasets used for the present study are available from the corresponding author upon reasonable request

