## Supplementary Figures for "Metabolic Stress Accelerates Dysregulated Synovial Macrophage-Fibroblast Communication and Htra1 Overproduction in Osteoarthritis"

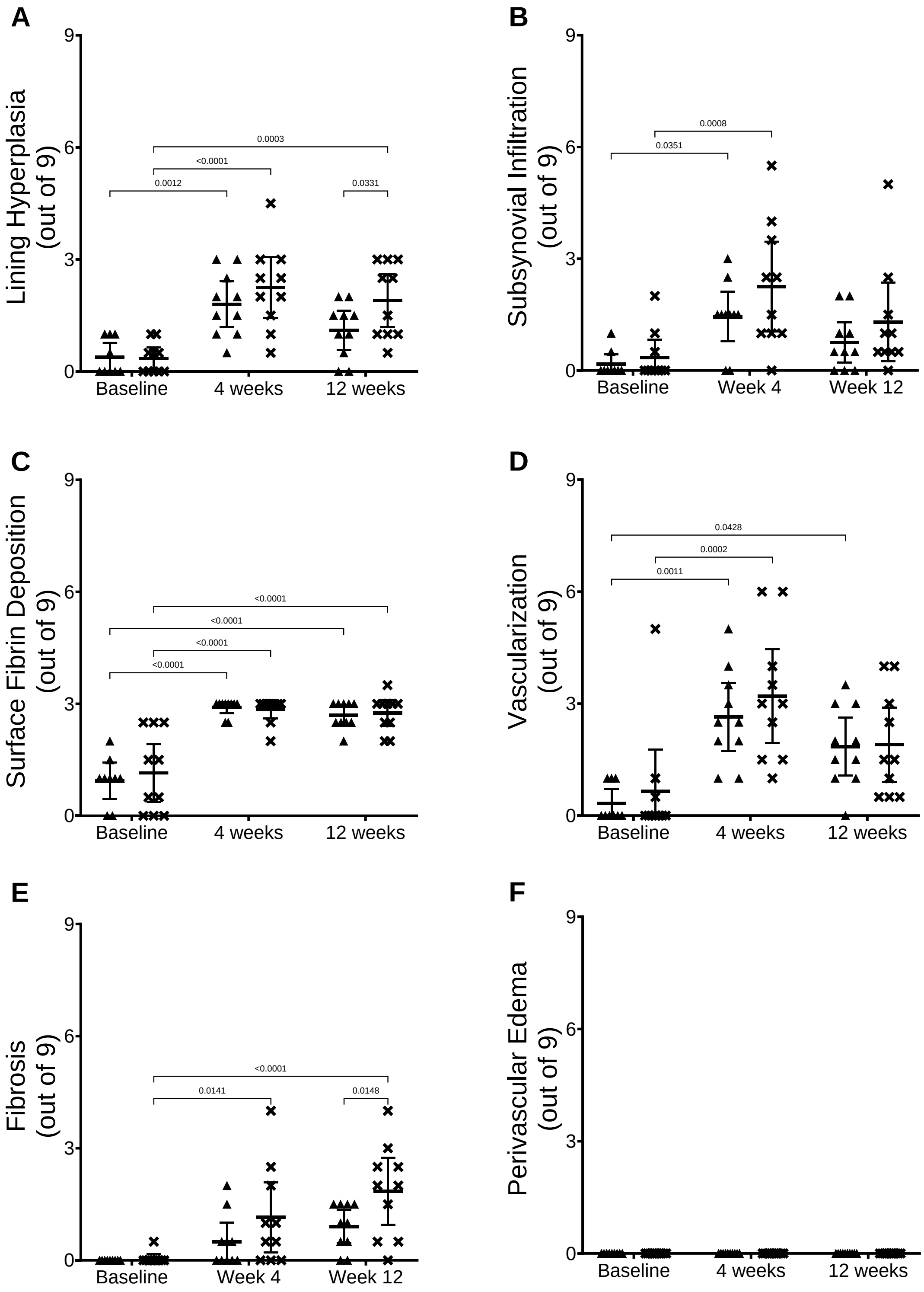


**Supplementary figure 1. Individual measures of synovial histopathology.** Related to figure 2. Measures of synovial histopathology in whole joint of RD (▲) or HFS (**x**) at baseline, 4-, and 12-week timepoints. The y-axis shows the total histopathological score (out of 9) for each component and the x-axis time. Mean with 95% confidence intervals are displayed.

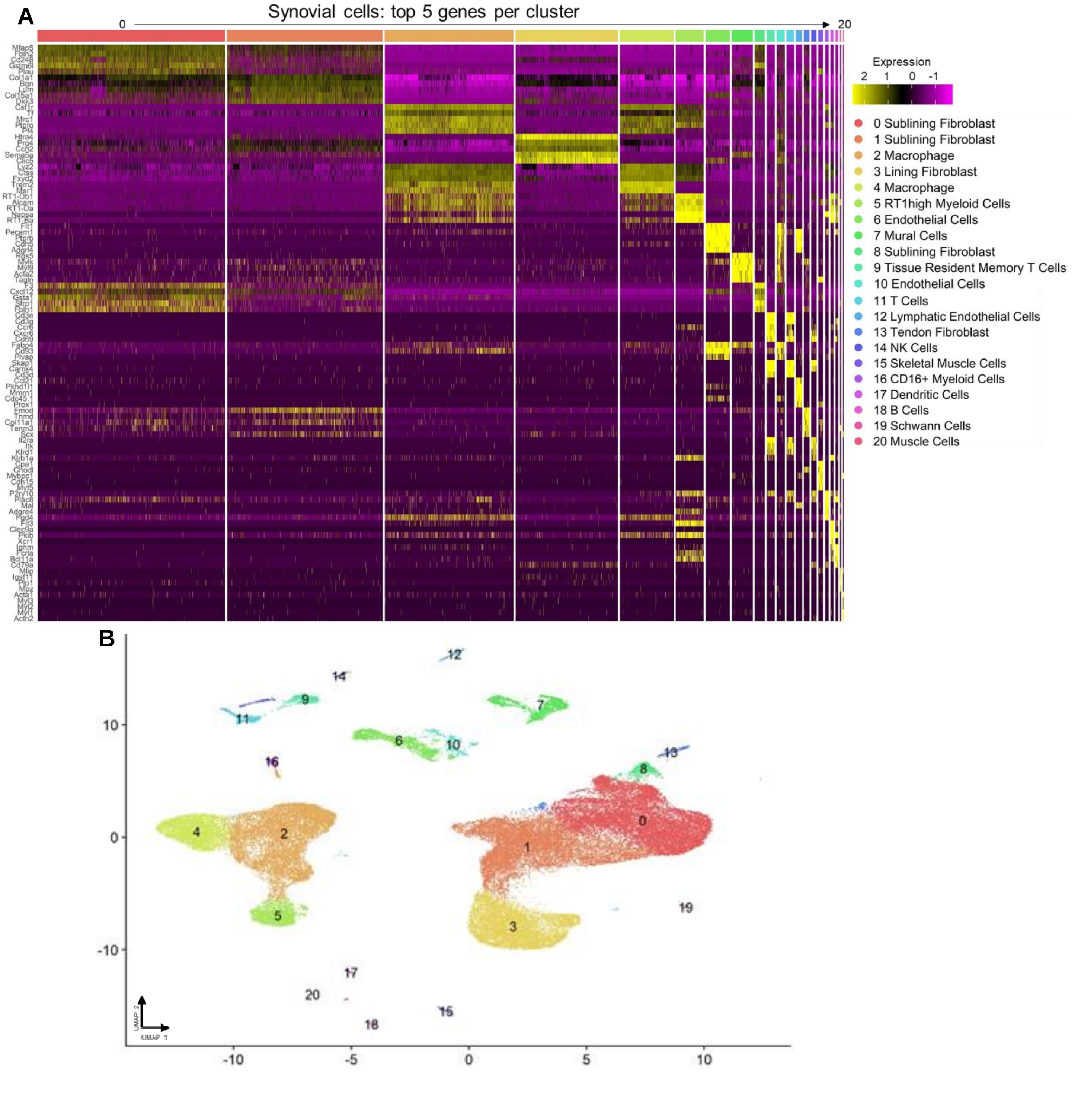


**Supplementary figure 2. Markers and annotations of all synovial cell types.** Related to figure 3. Heatmap showing the top 5 genes per clusters corresponding to the UMAP plots from figure 3A (**A**). UMAP showing all synovial cells from each condition (**B**).

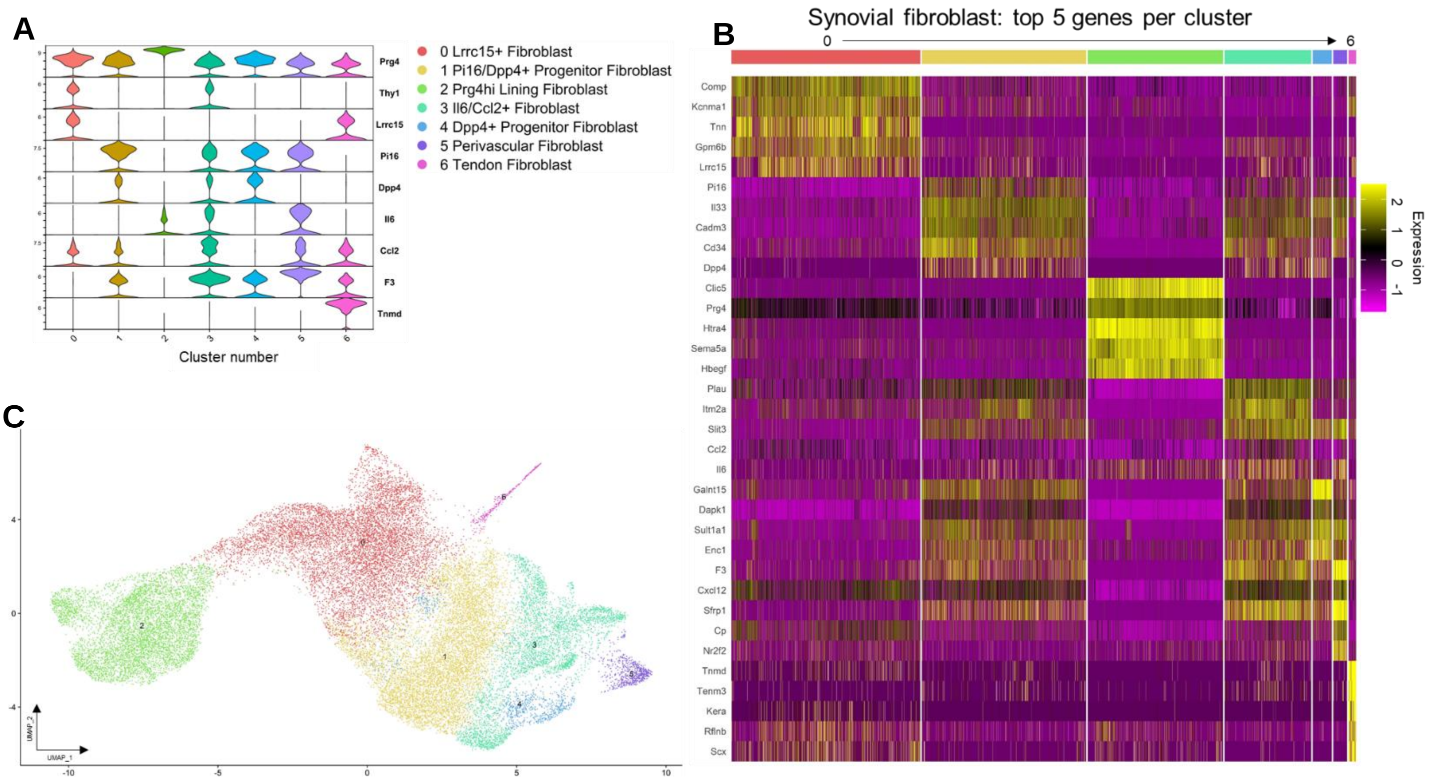


**Supplementary figure 3. Markers and annotations of synovial fibroblasts.** Related to figure 4. Violin plot showing individual markers of each cluster corresponding to the UMAP plots from figure 4A (**A**). Heatmap showing the top 5 genes per clusters corresponding to the UMAP plots from figure 4A (**B**). UMAP showing all synovial fibroblasts from each condition (**C**).

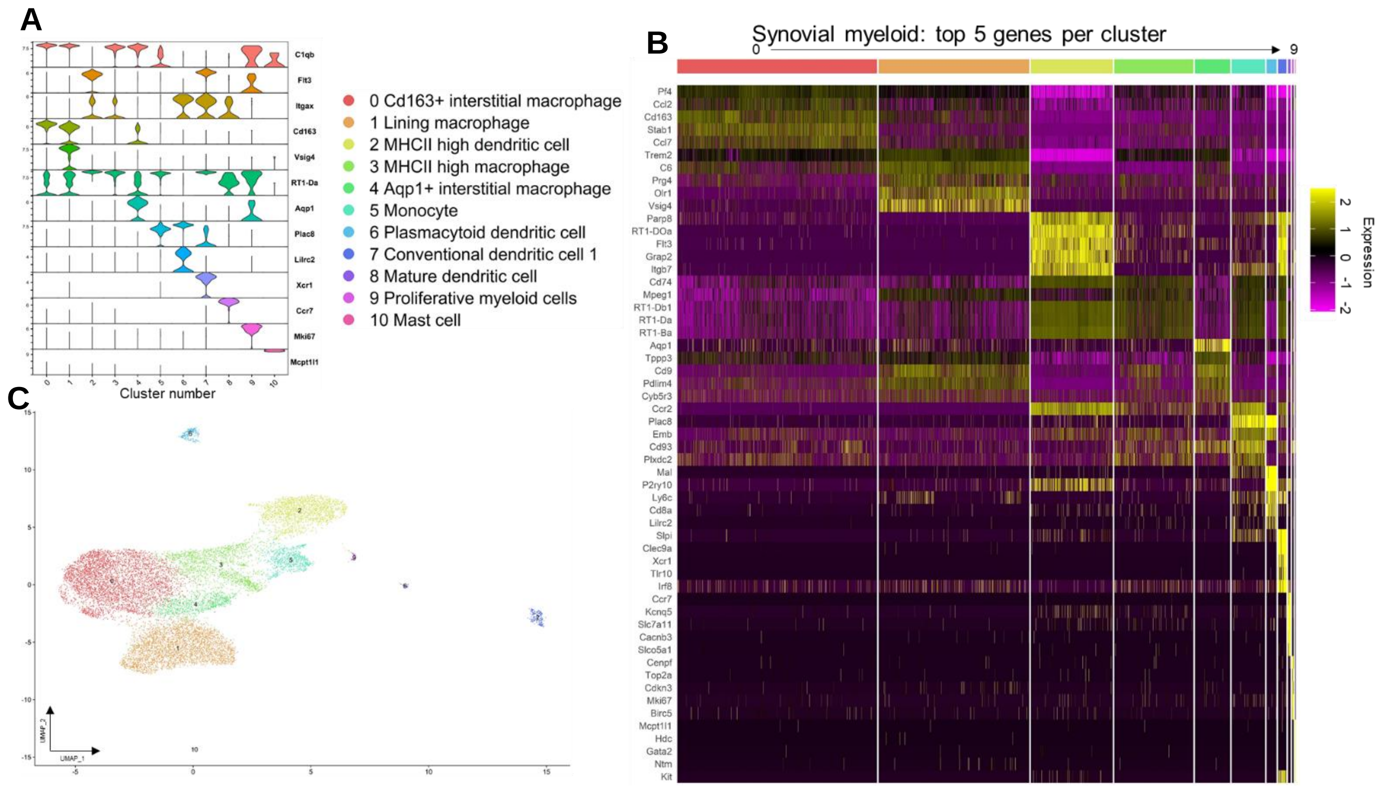


**Supplementary figure 4. Markers and annotations of synovial myeloid cells.** Related to figure 5. Violin plot showing individual markers of each cluster corresponding to the UMAP plots from figure 5A (**A**). Heatmap showing the top 5 genes per clusters corresponding to the UMAP plots from figure 5A (**B**). UMAP showing all synovial myeloid cells from each condition (**C**).
