## Supplementary Methods for "Metabolic Stress Accelerates Dysregulated Synovial Macrophage-Fibroblast Communication and Htra1 Overproduction in Osteoarthritis"

*Study participants*

Participants were recruited as part of the ongoing Western Ontario Registry for Early Osteoarthritis (WOREO) Knee Study, a prospective, single-centre, multi-clinic (St. Joseph’s Rheumatology clinic, Fowler Kennedy Sport Medicine Clinic, and Rorabeck Bourne Joint Replacement Clinic) cohort designed to investigate clinical, biomechanical, and pathophysiological features of early- and late-stage knee OA. Patients over the age of 18 with a diagnosis of early knee OA based on clinical assessment by a rheumatologist or orthopaedic surgeon and a Kellgren-Lawrence ≤2 were included. In patients with bilateral knee OA, the knee with worse symptoms was included. Patients with a history of inflammatory arthritis, disease-modifying anti-rheumatic drug (DMARD) use, oral corticosteroid use, or any knee procedure or corticosteroid injection within 6 months prior to enrollment assessment were excluded. Participants recruited consecutively between September 2017 - October 2023 were included in this analysis. Participants provided written informed consent and the registry was approved by Western University’s Research Ethics Board for Health Sciences Research Involving Human Subjects (HSREB #109255).

*Patient-reported measures of pain*

              Participants completed the KOOS Pain subscale (9 items: 0-100). Lower KOOS scores indicate worse pain. The KOOS questionnaire is valid and reliable for individuals with knee OA *(1, 2)*. A clinically meaningful difference in KOOS pain score is defined as a difference in 8.0-10.0 points *(3)*.

*Musculoskeletal ultrasound*

Knee ultrasound (US) scans were completed using a linear 3-12 MHz probe (GE LOGIQ, GE Healthcare) in accordance with OMERACT knee US protocol *(4)*. Each participant was supine with knees semi-flexed to 30-degrees. Longitudinal axis views were acquired in three standardized suprapatellar windows defined by the midline, lateral, and medial patellar poles. Synovitis severity was scored in each window from grade 0-3, according to the OMERACT knee US protocol *(4)*. Peak synovitis grade was assigned as the most severe score (0-3) of the three windows and used for these analyses. US scans were completed by 1 of 3 operators certified in musculoskeletal US by the Canadian Rheumatology Ultrasound Society who had at least 1-year musculoskeletal US experience.

*Rat model of experimental knee OA*

Twenty-four-week-old male Sprague-Dawley rats (Charles River Laboratory, Quebec, Canada, strain code 400) were housed and handled in the Animal Care and Veterinary Services facility at Western University in accordance with the guidelines of the Canadian Council on Animal Care. The animal use protocol was approved by the Western University Animal Care and Use Committee (AUP2021-087). Rats were fed either a control diet (RD) (3.0kcal/g, 12% fat and 2.7% sucrose) or a metabolic stress inducing high-fat-high-sucrose diet (HFS) (4.6kcal/g, 40% total energy from animal fat and 45% total energy from sucrose) (Dyets Inc., Bethlehem, PA, USA) for 12 weeks prior to OA induction or sacrificed for baseline (pre-OA, 0W) comparisons. Anterior cruciate ligament transection and destabilization of the medial meniscus surgery was used to induce experimental OA as previously described *(5)*. Rats were randomized to diet (RD or HFS) and OA timepoint (Baseline (0W), 4- (4W), or 12-weeks (12W).

*Pressure Application Measurement (PAM)*

Mechanical sensitivity at the knee was measured longitudinally in all animals in the 12-week endpoint group (n=14 per diet) as previously described *(6)*. Briefly, a pressure application measurement (PAM) algometer (Bioseb) was placed on the medial joint line of the knee and pressure was applied at approximately 200 grams per second until a withdrawal or vocalization occurred. Measurements were taken at baseline and 4-, 8-, and 12-week OA timepoints.

*Electronic von Frey (eVF)*

Using the same measurement approach as above, ipsilateral hindpaw mechanosensitivity was measured longitudinally in all 12-week endpoint rats as previously described *(6)*. In brief, force (25 grams per second) was applied to the plantar surface of the hindpaw by the applicator tip of an electronic von Frey instrument (Bioseb) until withdrawal occurred. Three measurements were taken with a 5-minute recovery period between.

*Cartilage and synovial histopathology*

Whole knees were harvested for histopathological analysis at baseline, 4-, and 12-week endpoints (n=9-10 per group), fixed, decalcified with 14% neutral-buffered EDTA, processed, sectioned, and stained with either toluidine blue or haematoxylin and eosin (H&E) as previously described *(6)*. Toluidine blue stained sections (3-5 per animal every ~600um to span the mid-tibiofemoral joint) were used for grading cartilage degeneration using the OARSI histopathology system *(7)*. Total joint degeneration scores were calculated by summing the mean grade of the medial and lateral femoral condyles and tibial plateaus. Synovial histopathology was assessed on the most anterior section (1 per animal) of an H&E stained slide using the six-component scoring system *(8)*. Mean grades for the parapatellar and tibiofemoral regions were collected for synovial lining thickness, synovial infiltration, fibrin deposition, vascularization, fibrosis, and perivascular edema. Mean grades from each anatomical region were summed for a total joint score for each component. The average score of two blinded expert readers was used for both cartilage and synovitis grading.

*Synovial tissue dissociation and preparation of single cell suspension*

Synovial tissue (n=2-3 per condition) was dissected away from underlying connective tissue, rinsed in phosphate buffered saline (PBS), minced into small pieces, and enzymatically digested as previously described *(6)*. In brief, tissue pieces were digested in complete Roswell Park Memorial Institute media (RPMI) (10% fetal bovine serum, 1% penicillin/streptomycin) containing 400 µg/mL Liberase^TM^ and 400 µg/mL DNase I. Next, 100 mM EDTA was added to stop the enzymatic reaction and cells were washed twice with Dulbecco’s Modified Eagle Medium (DMEM) prior to cell counts and live/dead quantification. Cells were aliquoted into 1-2 million cells/mL in CryoStor® CS10 cryopreservation media, frozen at -80°C overnight in a controlled-rate cell freezing container and transferred to liquid nitrogen vapor phase storage until use.

*Single cell RNA library preparation and sequencing*

Synovial single cell suspensions were rapidly thawed from cryopreservation. Cell suspensions were washed then counted and viability assessed with trypan blue exclusion. Samples with a viability of ≥ 62% were chosen for library preparation.

Synovial cells were prepared for cDNA amplification and chromium library construction using the 10x Chromium Next GEM Single Cell 3' Library and Gel Bead Kit v3.1 according to manufacturer’s protocol (10x Genomics, Pleasanton, CA, USA) at the Lawson Single Cell Barcoding Facility (London, ON, Canada). The cDNA libraries were sequenced on the Illumina NovaSeq6000, using the S4 flow cell with a read length of 100 bp (paired end) by the Genome Quebec core facility (Montreal, QC, Canada). The average read depth was approximately 59,376 reads per cell.

*Analysis of single cell sequencing data*

Sequencer files were decompressed into FASTQ format prior to alignment. Raw data was aligned to the mRatBN7 (1.0.0) reference rat transcriptome using STAR. Filtering, barcoding, and UMI counting were performed using Cell Ranger 7.0.0 and the 10X Genomics recommended default parameters. All downstream analysis was performed in Rstudio (v 4.2.2) using the Seurat (v4.3.1) package. Cells with less than 100 genes and genes expressed in fewer than 10 cells were excluded. Low-quality cells were removed on a per sample basis based on gene count (nCount) and number of expressed genes (nFeature). Remaining cells had 5000 < nCount < 93^rd^ percentile and 10^th^ percentile < nFeature < 98^th^ percentile. Cells were excluded if mitochondrial-derived genes exceeded 10% of all genes. DoubletFinder (v2.0.3) was applied to identify and exclude doublets *(9)*. Harmony integration was trialed but did not influence the dataset and was not incorporated. Counts were normalized using *LogNormalize* with a scale factor of 1,000,000. The top 2000 variable genes were identified using *FindVariableFeatures* and VST. Principal component analysis was performed, and elbow plots used to determine the optimal number of dimensions (35 PCs) for subsequent dimensionality reduction by Uniform Manifold Approximation and Projection (UMAP). Unsupervised clustering was performed using *FindNeighbors* and *FindClusters,* resolution = 0.2.

Cluster specific metrics were identified using *FindAllMarkers* with a log2 fold change threshold of 0.5. Clusters were manually annotated based on previously published synovial single cell datasets *(10–15)*. Cell type specific analysis was performed to identify myeloid and fibroblast subsets. In short, fibroblasts (not including pericytes) or myeloid cells were subset from the “all cells” object and re-clustered (fibroblasts; 40 PC’s and resolution 0.2, or myeloid; 40 PCs and resolution 0.3). Cluster annotations were performed as described above. Functional annotation was performed by submitting the genes output by *FindAllMarkers* to Gene Ontology (GO) pathway analysis (Biological Processes) using PantherDB. To identify global gene expression differences of all cells, myeloid cells, or fibroblasts a pseudobulk method and DESeq2 were used. For gene set enrichment, the differentially expressed genes (adjusted p-value < 0.05) and their log2 fold change were submitted to GO Biological Processes (GO:BP) and Reactome using PantherDB. Bubble plots for enriched terms and functional annotations were generated with ggplot2.

Crosstalk between synovial macrophages and fibroblasts was assessed using CellChat (v1.6.1) *(16)*. All analysis was performed using the provided mouse database. Over-expressed genes and ligand-receptor interactions were identified with *identifyOverExpressedGenes* and *identifyOverExpressedInteractions*. Data was projected against the provided PPI.mouse dataset to reduce effects of drop out. Cell-cell communication probabilities were calculated using *computeCommunProb* and *triMean* with a minimum cell number of 10 per cluster, and a significance threshold of 0.05. Pattern numbers for river plots were chosen when cophenetic and silhouette measures decreased simultaneously. Significantly communicating macrophage-to-fibroblast pathways were ranked by ratio of probabilities relative to all communication probabilities and pathways with probabilities greater than 1 were compared to identify unique communication pathways in each condition compared to regular diet.

*Immunofluorescence*

Immunofluorescence labeling was used to quantify 3-nitrotyrosine-positive (3NT) synoviocytes in rat joint sections (n = 3 per group) *(17)*. Sections were labeled with rabbit anti-3NT (Millipore, 0.02 mg/mL) or rabbit polyclonal isotype control (Abcam, ab37415) in blocking buffer overnight at 4^o^C. Then, goat anti-rabbit Alexa 488 secondary antibody (Jackson ImmunoResearch, 0.0015 mg/mL) was applied to all sections before mounting with ProLong Gold Antifade Mountant with DAPI (Fisher Scientific). All slides were imaged on a Leica SP8 confocal microscope using the 40x oil-immersion objective with constant settings across all samples. Two fields of view were selected and imaged per section.

*Macrophage-fibroblast co-culture*

Peripheral blood mononuclear cells were isolated from a single healthy donor as previously described *(17)*. Following isolation, 125000 mononuclear cells were seeded per well and cultured for seven days in either normal glucose (11.1mM) (NGlu) or high glucose (31.1mM) (HGlu) complete RPMI containing 50ng/mL macrophage colony-stimulating factor (mscf) to prepare healthy or metabolically stressed macrophages *(18)*. On day seven, cultures were washed providing highly enriched blood-derived macrophages for co-culture. Additional NGlu and HGlu wells (n=3) were prepared as above and cultured in NGlu complete RPMI with 50ng/mL mcsf for co-culture controls.

OA synovial fibroblasts (OASF) were isolated from synovial tissue donated by WOREO patients at the time of joint arthroplasty or high tibial osteotomy. Synovial tissue (n=6) was dissociated as described above and OASF were enriched through passage. One hundred thousand passage 4 OASF were seeded onto washed NGlu or HGlu macrophages and cultured for 72 hours in normal glucose complete RPMI containing 50ng/mL mcsf. HTRA1 was measured in conditioned media from co-culture or macrophage controls by ELISA (My BioSource) per the manufacturer’s specifications. All samples were tested in duplicate.
