## Supplementary Tables 1-3 for "Metabolic Stress Accelerates Dysregulated Synovial Macrophage-Fibroblast Communication and Htra1 Overproduction in Osteoarthritis"

| **Supplementary Table 1. Multivariable linear regression evaluating the association between metabolic syndrome with KOOS pain scores** | | | |
| --- | --- | --- | --- |
| **Primary analysis** |  |  |  |
| **Variable** | **β coefficient** | **Standard Errors** | **95% CIs** |
| **MetS** |  |  |  |
| No | Reference | Reference | Reference |
| Yes | **-10.90** | **3.14** | **-17.11 to -4.70** |
| **Including US-synovitis as covariate*** | |  |  |
| **Variable** | **β coefficient** | **Standard Errors** | **95% CIs** |
| **MetS** |  |  |  |
| No | Reference | Reference | Reference |
| **Yes** | **-9.70** | **3.26** | **-16.15 to -3.26** |
| Adjusting for age and sex only  *Adjusting for age, sex, and US-synovitis (none/mild vs moderate/severe)  Values in bold indicate significance at the 5% level  CI, confidence interval; KL, Kellgren-Lawrence grade; KOOS, Knee Injury and Osteoarthritis Outcome Score; MetS, metabolic syndrome; US, ultrasound. | | | |

| **Supplementary table 2. Linear mixed-effects model estimates of pressure application measurements for RD (A) and HFS (B) and pairwise comparisons (C).** | | | |
| --- | --- | --- | --- |
| **A** | | |  |
| ***Fixed Effects*** | | |  |
| **Variable** | **β coefficient** | **Std Err** | **95% CI** |
| **Time** |  |  |  |
| Baseline | Reference | Reference | Reference |
| **Week 4** | **-68.73** | **31.97** | **(-130.52, -6.94)** |
| Week 8 | 56.62 | 31.97 | (-5.17, 118.41) |
| Week 12 | 16.49 | 31.97 | (-45.30, 78.27) |
| **Intercept** | 632.49 | 23.3 | (587.71, 677.26) |
| ***Random Effects*** |  |  |  |
| **Variable** | **Variance** | **Std Dev** |  |
| Intercept (Animal ID) | 444.4 | 21.08 |  |
| Residual | 7154.5 | 84.58 |  |

**B**

| ***Fixed Effects*** | | |  |
| --- | --- | --- | --- |
| **Variable** | **β coefficient** | **Std Err** | **95% CI** |
| **Time** |  |  |  |
| Baseline | Reference | Reference | Reference |
| **Week 4** | **-243.26** | **28.02** | **(-297.36, -189.17)** |
| **Week 8** | **-121.05** | **28.56** | **(-176.04, -65.78)** |
| **Week 12** | **-157.95** | **28.56** | **(-212.94, -102.68)** |
| **Intercept** | 760.16 | 19.99 | (721.78, 798.54) |
| ***Random Effects*** |  |  |  |
| **Variable** | **Variance** | **Std Dev** |  |
| Intercept (Animal ID) | 98.05 | 9.9 |  |
| Residual | 5495.25 | 74.13 |  |

**C**

| **Diet x Time** | **Estimate** | **Std Err** | **95% CI** |
| --- | --- | --- | --- |
| **Baseline** | **127.67** | **30.7** | **(32.52, 222.82)** |
| Week 4 | -46.86 | 30.7 | (-142.02, 48.29) |
| Week 8 | -50.19 | 31.3 | (-147.19, 46.82) |
| Week 12 | -46.95 | 31.3 | (-143.96, 50.05) |

RD: regular diet; HFS: high-fat-high-sucrose diet; Std Err: standard error; 95%CI: 95% confidence interval. Adjusted for animal identification (Animal ID). Values in bold indicate significance at the 5% level.

| **Supplementary table 3. Linear mixed-effects model estimates of electronic Von Frey measurements for RD (A), HFS (B), pairwise comparisons (C).** | | | |
| --- | --- | --- | --- |
| **A** | | |  |
| ***Fixed Effects*** | | |  |
| **Variable** | **β coefficient** | **Std Err** | **95% CI** |
| **Time** |  |  |  |
| Baseline | Reference | Reference | Reference |
| **Week 4** | **-14.61** | **3.74** | **(-21.83, -7.39)** |
| **Week 8** | **-10.96** | **3.74** | **(-18.18, -3.74)** |
| **Week 12** | **-18.26** | **3.74** | **(-25.48, -11.04)** |
| **Intercept** | 84.88 | 2.82 | (79.46, 90.29) |
| ***Random Effects*** |  |  |  |
| **Variable** | **Variance** | **Std Dev** |  |
| Intercept (Animal ID) | 13.31 | 3.65 |  |
| Residual | 97.7 | 9.88 |  |

**B**

| ***Fixed Effects*** | | |  |
| --- | --- | --- | --- |
| **Variable** | **β coefficient** | **Std Err** | **95% CI** |
| **Time** |  |  |  |
| Baseline | Reference | Reference | Reference |
| **Week 4** | **-9.43** | **3.45** | **(-16.09, -2.78)** |
| **Week 8** | **-8.93** | **3.53** | **(-15.76, -2.11)** |
| **Week 12** | **-7.32** | **3.53** | **(-14.14, -0.50)** |
| **Intercept** | 83.21 | 3.42 | (76.54, 89.88) |
| ***Random Effects*** |  |  |  |
| **Variable** | **Variance** | **Std Dev** |  |
| Intercept (Animal ID) | 80.99 | 9 |  |
| Residual | 83.17 | 9.12 |  |

**C**

| **Diet x Time** | **Estimate** | **Std Err** | **95% CI** |
| --- | --- | --- | --- |
| **Baseline** | -1.67 | 4.43 | (-15.49, 12.16) |
| Week 4 | 3.51 | 4.43 | (-10.31, 17.33) |
| Week 8 | 0.36 | 4.51 | (-13.68, 14.40) |
| Week 12 | 9.28 | 4.51 | (-4.76, 23.31) |

RD: regular diet; HFS: high-fat-high-sucrose diet; Std Err: standard error; 95%CI: 95% confidence interval. Adjusted for animal identification (Animal ID). Values in bold indicate significance at the 5% level.
